# Epimutations driven by small RNAs arise frequently but have limited duration in a metazoan organism

**DOI:** 10.1101/2019.12.29.890194

**Authors:** Toni Beltran, Vahid Shahrezaei, Vaishali Katju, Peter Sarkies

## Abstract

Epigenetic regulation involves changes in gene expression independent of DNA sequence variation that are inherited through cell division (Holliday, 2006). In addition to a fundamental role in cell differentiation, some epigenetic changes can also be transmitted transgenerationally through meiosis (Heard and Martienssen, 2014). Epigenetic alterations (“epimutations”) could thus contribute to heritable variation within populations and be subject to evolutionary processes such as natural selection and drift (Burggren, 2016). However, this suggestion is controversial, partly because unlike classical mutations involving DNA sequence changes, key parameters such as the rate at which epimutations arise and their persistence are unknown. Here, we perform the first genome-wide study of epimutations in a metazoan organism. We use experimental evolution to characterise the rate, spectrum and stability of epimutations driven by small silencing RNAs in the model nematode *C. elegans*. We show that epimutations arise spontaneously at a rate ∼25 times greater than DNA sequence changes and typically have short half-lives of 2-3 generations. Nevertheless, some epimutations last at least 10 generations. Epimutations thus may contribute to evolutionary processes over a short timescale but are unlikely to bring about long-term divergence without further DNA sequence changes.

---

In the nematode *Caenorhabditis elegans*, an epigenetic memory of gene silencing can be transmitted transgenerationally for multiple generations. This extremely stable form of silencing is initiated by Piwi-interacting small RNAs (piRNAs), leading to the formation of secondary small RNAs known as 22G-RNAs by RNA-dependent RNA polymerase (RdRP) activity (Ashe et al., 2012; Luteijn et al., 2012; Shirayama et al., 2012). 22G-RNAs and their associated Argonaute HRDE-1 are transmitted through gametes, and are required for the maintenance of silencing each generation (Buckley et al., 2012), where 22G-RNA amplification can continue independently of the initial trigger (Sapetschnig et al., 2015).

So far, mechanistic investigation of transgenerational epigenetic inheritance of 22G-RNAs has largely been confined to transgenes and artificial induction of RNAi from exogenous sources (Ashe et al., 2012; Buckley et al., 2012; Luteijn et al., 2012; Shirayama et al., 2012). The transgenerational dynamics of small RNA populations targeting endogenous genes are, in contrast, very poorly understood (de Albuquerque et al., 2015; Phillips et al., 2015). In particular, whether epigenetic inheritance can provide an additional layer of heritable biological variation for evolutionary forces to act on remains obscure. This idea is controversial, since key parameters regarding the stability of epigenetic states across generations are unknown.

To investigate whether 22G-RNA-mediated epimutations occur at endogenous genes in *C. elegans*, we used a mutation accumulation (MA) approach. Mutation accumulation lines are used in classical evolutionary biology in order to provide unbiased estimates of mutation rates (Katju and Bergthorsson, 2019). In the MA approach, multiple lines are propagated independently from a common ancestor for several generations. Crucially, the population is repeatedly passed through bottlenecks containing very few individual organisms. Since *C. elegans* is a hermaphrodite, the minimal bottleneck size is a single individual. This approach strongly reduces the ability of natural selection to eliminate deleterious mutations, thus enabling an unbiased estimate of the true spectrum of mutations to be made (Katju and Bergthorsson, 2019). We employed *C. elegans* MA lines grown at a population size of 1 over multiple generations (Katju et al., 2018; Konrad et al., 2018) to investigate epimutations (Fig. 1a). We reasoned that loci subject to epimutations would show alterations in 22G-RNA levels relative to the parental line. Such changes could arise in individual lines as a result of stochastic processes such as environmental fluctuations, developmental noise, or intrinsic noise in the pathways of epigenetic regulation. Crucially, since the lines were maintained under identical conditions (Katju et al., 2018; Konrad et al., 2018), long term maintenance of 22G-RNA alterations within lines over multiple generations might indicate epigenetic transmission.

**Figure 1.**
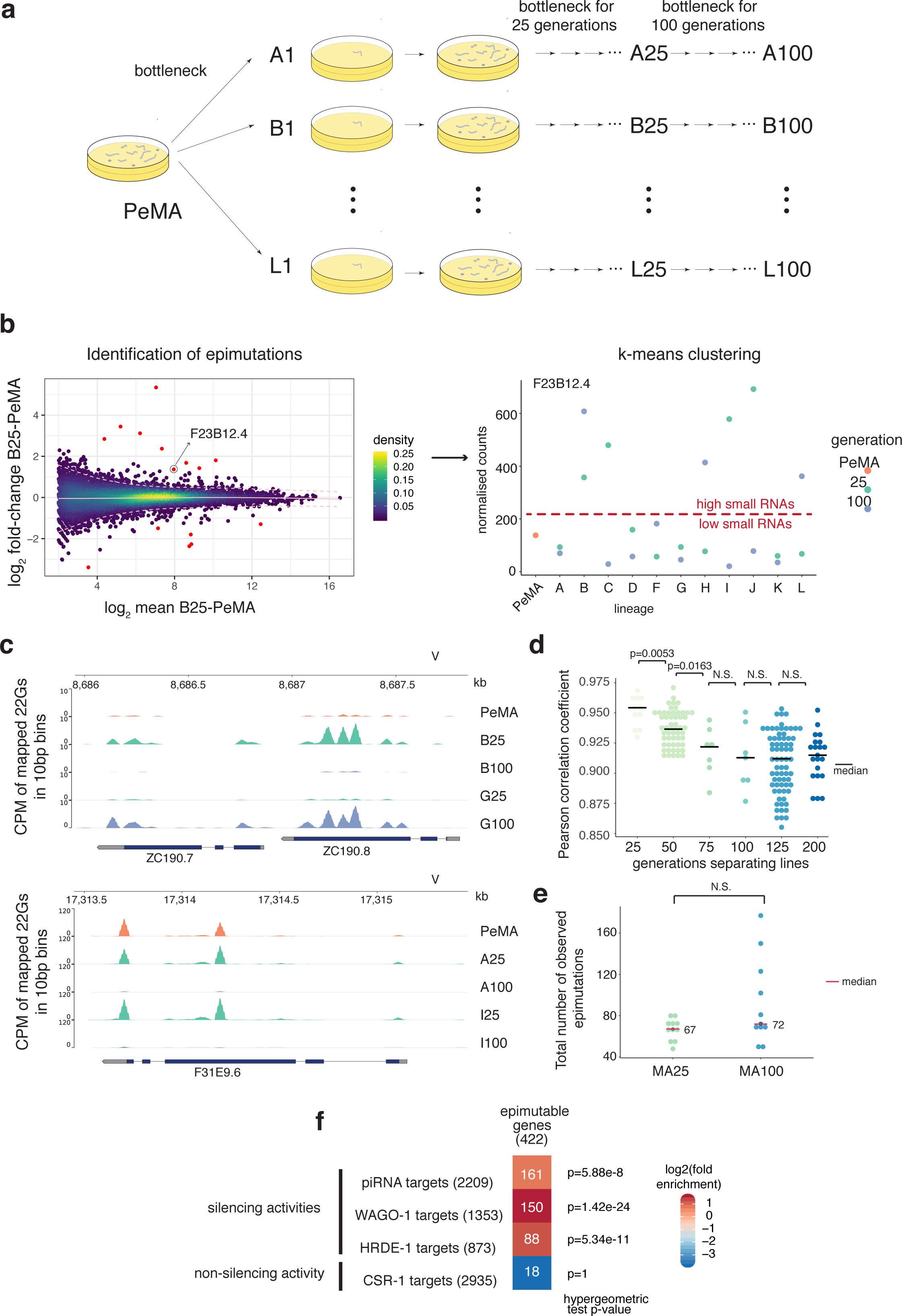
Identification of small RNA-mediated epimutations in *C. elegans* mutation accumulation lines. a. Experimental design. 11 lineages of *C. elegans* nematodes were grown at a population size of 1 individual, for 100 generations. Small RNA sequencing was carried out for each lineage at 25 and 100 generations, as well as the parental strain before epimutation accumulation (PeMA). b. Identification of epimutations. Epimutable genes are defined as genes with large fold changes in 22G levels in at least one pairwise comparison, compared to genes of similar 22G abundance (FDR<1e-4). K-means clustering was applied on the normalised count data to define groups of samples with high and low small RNA levels. c. 22G signal profiles for representative examples of epimutations. The top panel shows an epimutation affecting two nearby genes. d. Dot plots depicting the distribution of Pearson correlation coefficients between lines as a function of the distance in number of generations. Significance was tested using a two-sided Wilcoxon rank sum test. e. Number of epimutations detected after 25 and 100 generations. f. Overlap of epimutated genes with previously annotated small RNA pathway target genes. The significance of the overlap is indicated by the colour of the heatmap.

We used high throughput small RNA sequencing to map candidate epimutations in 11 MA lines after 25 and 100 MA generations (MA25 and MA100 respectively). We were able to robustly identify genes with significantly larger changes in 22G-RNA levels than expected given their mean 22G-RNA level (FDR<10^−4^, Methods; examples in Fig. 1b and Fig. 1c). We detected a total of 422 genes with significant changes in 22G-RNAs in at least one MA line. These events are candidates for epimutations as they may be inherited across generations. On average, we detected around 70 candidate epimutations in each line, and the overall number of candidate epimutations was similar in MA25 and MA100 lines (Fig. 1d,e, Supp. Fig. 1). Notably, over 25 generations, the expected number of fixed point mutations and small indels is ∼4.61 according to the estimated mutation rate from the same lines (Konrad et al., 2019). In addition, only a tiny fraction of mutations detected after 400 generations (Konrad et al., 2019) overlapped with epimutations (Supp. Fig. 2a-c). The majority of changes in 22G-RNAs are therefore unlikely to be due to genetic differences.

Genes subject to changes in 22G-RNAs overlapped significantly with piRNA (Tang et al., 2016), HRDE-1 (Shirayama et al., 2012) and WAGO-1 (Gu et al., 2009) target genes, but not with genes targeted by CSR-1 (Claycomb et al., 2009), which has been proposed to oppose the silencing activity of other germline RNAi pathways (Seth et al., 2013; Wedeles et al., 2013) (Fig. 1f). We also detected several changes in 22G-RNAs at transposable elements (TEs), and indeed TEs were enriched for these alterations (Supp. Fig. 9a,b). Together, this suggests that stochastic changes in small RNAs at individual genes and repetitive elements occur frequently during *C. elegans* propagation under minimal population size.

Changes in small RNAs in individual lines could reflect changes driven by environmental or developmental fluctuations lasting only one generation or could reflect epigenetic inheritance of changes that occurred in previous generations. To test whether long-term inheritance of changes in small RNAs occurs, we compared individual genes between MA25 and MA100 lines. Only a minority of changes in 22G-RNA levels were retained (13±3% mean and 95% confidence intervals; Fig. 2a, Supp. Fig. 9c). Given the fast rate at which changes in small RNAs arose, the small percentage shared between the same MA25 and MA100 line may have arisen independently. Indeed, across the set of genes, alterations in 22G-RNAs were no more likely to be shared between the same MA25 and MA100 line than expected by chance (Fig. 2b,c). Similarly, there was no significant tendency for the variance in 22G-RNA levels at either genes or TEs to be lower within lineages than between lineages (Fig. 2b,c; Supp. Fig. 9d). We therefore conclude that the alterations in 22G-RNA levels have limited stability such that few survived over the course of 75 generations (examples in Fig. 2d,e; Supp. Fig. 3a,b for the few examples reaching significance).

**Figure 2.**
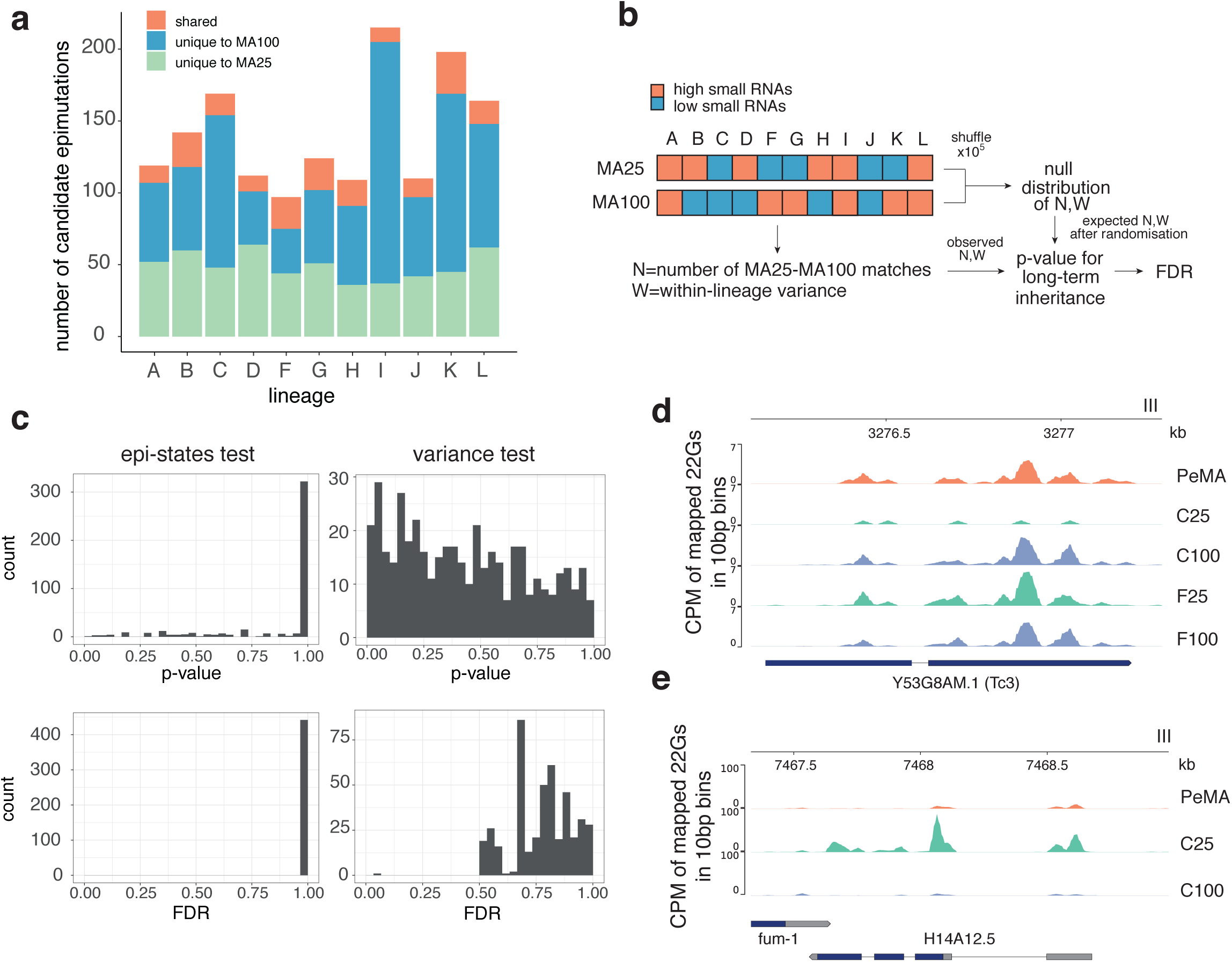
Absence of evidence for long-term inheritance of epimutations. a. Unique and shared epimutation totals in MA25 and MA100 lines, showing that only a small fraction of epimutations is maintained. b. Test for long-term epigenetic inheritance. The number of matching states in pairs of MA25 and MA100 lines of the same lineage was calculated, and compared to the expected number of matches when randomly pairing MA25 and MA100 lines (epi-states test; see methods). Similarly, the variance in pairs of MA25 and MA100 lines of the same lineage was calculated and compared to the expected variance in random pairs (variance test; see methods). c. Histograms of p-values and Benjamini-Hochberg False Discovery Rate-corrected p-values of both tests, showing that virtually no cases remain significant after multiple testing correction. d-e. Genome browser windows showing examples of unstable epimutations mapped onto the *C. elegans* genome (ce11).

Having established that long-lasting epimutations are rare in *C. elegans*, we sought to investigate the persistence of alterations in 22G-RNAs over shorter timescales. We propagated two MA lines for 13 generations, and sequenced small RNAs at each generation (Fig. 3a). We identified genes with significant (FDR<10^−4^) changes in 22G-RNA levels between any two generations within each lineage. The genes identified in this experiment overlapped significantly (35%) with the candidate epimutations in MA25 and MA100 lines, and with piRNA, HRDE-1 and WAGO-1 targets, but not CSR-1 targets (Fig. 3b). Within this set of genes, the correlation coefficients between samples decreased with increasing distance in generations, suggesting that the level of 22G-RNAs targeting these genes is inherited transgenerationally (Fig. 3c).

**Figure 3.**
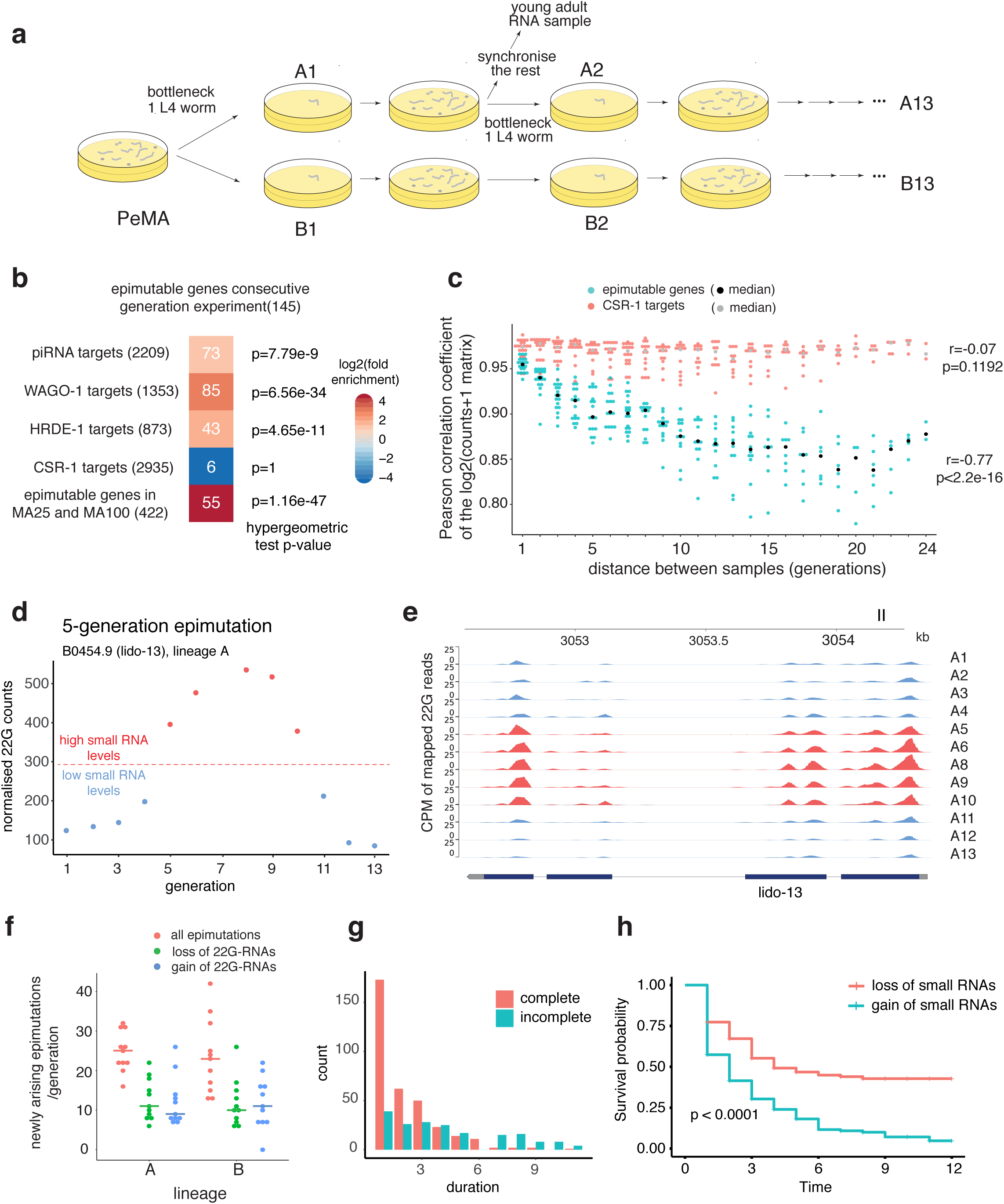
Characterisation of the short term dynamics of epimutations. a. Experimental design. Two 13-generation lineages were grown, by picking a single L4 stage worm as a founder for the next generation. For each generation, the remaining worm populations were synchronised using hypochlorite treatment and grown to adulthood to obtain RNA. b. Overlap of epimutated genes with small RNA pathway target genes. c. Correlation coefficients between samples in the consecutive generation experiment, as a function of separation in number of generations. d-e. Example of an epimutation lasting 5 generations. f. Dot plot showing the distribution of the number of newly arising epimutations in each of the generations of lineages A and B. g. Distribution of epimutation duration. Epimutations were split in two groups according to whether they revert within the lineage (complete), or whether they remain stable until the end of the observation window (incomplete, censored). h. Survival curves for epimutations consisting of a gain or a loss of small RNAs.

To characterise the rate and stability of alterations in 22G-RNAs we classified the count data across the two lineages into high and low small RNA level states (Methods, Fig. 3d, Fig. 3e). Based on this classification, we observed a median of 23.5 newly arising changes in 22G-RNAs per generation (Fig. 3f). This figure can be compared directly with the rate of DNA sequence mutations. Single nucleotide substitution and small indel rate estimates range from 0.2 to 1 changes per genome per generation (Denver et al., 2004, 2009; Konrad et al., 2019; Meier et al., 2014). Similarly, the rate of larger genomic duplications and deletions has been estimated to be of 6.5×10^−3^ per genome per generation (Konrad et al., 2018). The rate at which changes in 22G-RNAs arise is therefore at least 25 times greater.

Crucially, Many changes in 22G-RNAs persist for several generations, marking them out as genuine epimutations that can be transmitted transgenerationally (examples in Fig. 3d, Fig. 3e, Fig. 3g). We quantified the duration of epimutations using survival analysis, revealing a median survival of 2-3 generations considering all epimutations (Fig. 3h). Epimutations exhibiting increased 22G-RNAs (gains) had a median survival of 2 generations, while epimutations with decreased 22G-RNAs (losses) had a significantly greater median survival of 4 generations (*p*<1e-4 Mantel log-rank test, Fig. 3h). Interestingly, 42% of losses of 22G-RNAs remained stable after 12 generations, in contrast to only 5% of gains of 22G-RNAs (Fig. 3h). Gains of 22G-RNAs affecting genes targeted by piRNAs, WAGO-1 or HRDE-1 tended to have increased stability, while the few occurring in CSR-1 targets were extremely unstable (Supp. Fig. 4), consistent with a protective role of CSR-1 against stochastic silencing(Seth et al., 2013; Wedeles et al., 2013).

Interestingly, some epimutations showed a gradual change in 22G-RNAs (Fig. 3d), while others showed a clear shift between high and low small RNAs, indicating the possibility of bistability (Supp. Fig. 5a). To examine this across all epimutations, we examined the change in 22G-RNA levels across a 6-generation window centred on the generation in which the epimutation was identified (Supp. Fig. 5b,c). We identified three categories of epimutation. Most genes showed either fluctuating small RNA levels or evidence of bistable behaviour, while less than 10% (8/145) showed evidence of a gradual change in 22G-RNAs (Supp. Fig. 5d,e). Epimutable genes showing bistable and gradual changes were significantly enriched for piRNA targets (p<0.01, Fisher’s Exact Test), but not for HRDE-1 or WAGO-1 targets (Supp. Fig. 5f). Bistable and gradual changes tended to last longer than epimutations at genes displaying fluctuating levels of 22G-RNAs (Supp. Fig. 5g).

The existence of different types of transgenerational dynamics in 22G-RNA levels prompted us to test the heritability of small RNA levels using an alternative method. We reasoned that transgenerational inheritance of 22G-RNA levels would result in reduced variance between consecutive generations (“intergenerational variance”) compared to the total variance (Supp. Fig. 6a). We identified 321 genes following this premise (permutation test FDR<0.1; Supp. Fig. 6b,c). Within these genes, we extracted the duration of runs of consecutive generations with low intergenerational variance (Supp. Fig. 6d), recovering a median duration of 2 generations. This analysis thus further supports the existence of heritable variation in 22G-RNA levels that lasts for ∼2-3 generations on average (Supp. Fig. 6e).

To examine the effect of small RNA-mediated epimutations on gene expression, we performed high-throughput RNA sequencing across the 12 generations, as well as MA25 and MA100 lines. Genes exhibiting epimutations were enriched for correlated changes in mRNA levels (Supp. Fig. 7a,c; simulation test p=0.036), and larger absolute changes in 22G-RNA levels were correlated with larger changes in mRNA levels (Supp. Fig. 7b). Changes in mRNA levels in the opposite direction to changes in 22G-RNAs were enriched (simulation test p=0.01, Supp. Fig 7c) but we also observed changes in the same direction, indicating that epimutations sometimes, but not always, correspond to gene silencing (Supp. Fig. 7d,e). It is possible that correlated increases in both small RNA and mRNA levels represent small RNA-mediated compensation for changes in expression level driven by other epigenetic changes. We find that the proportion of this class of epimutations is significantly higher in repressive chromatin regions (Evans et al., 2016) (Supp. Fig. 7f,g), suggesting that small RNAs might be responding to a loss of chromatin-mediated silencing.

Given the limited duration of epimutations, we speculated that the epimutations that we identify might represent extreme examples of fluctuations in some classes of small RNA levels at a subset of genes. We used our sequencing data to estimate the variability in 22G-RNAs across lineages, and compared this value to an estimate of the technical noise (Fig. 4a). 22G-RNAs levels are clearly more variable than technical noise, validating our approach, and allowing us to define genes with hypervariable small RNAs (“HV22Gs”) (FDR<0.1 in MA25, FDR<0.01 in MA100, Fig. 4a). No miRNAs and very few piRNAs showed increased variability, demonstrating that this effect is specific for 22G-RNAs (Supp. Fig 8a,b). HRDE-1, WAGO-1, piRNA target genes and transposable elements showed higher variability in small RNA levels and were enriched for HV22Gs, while CSR-1 targets were depleted (Fig. 4b-f; Supp Fig. 9e,f). Almost all epimutable genes were recovered as HV22Gs in this analysis (Fig. 4c). Genes identified as epimutated exclusively in MA25 lines, but not in MA100 lines nevertheless showed increased variability in the MA100 lines, and vice versa (Supp. Fig. 8c,d). This supports the hypothesis that epimutable genes show hypervariability in 22G-RNAs. HV22Gs did not show increased variability at the mRNA level (Supp. Fig. 8e,f). Interestingly, however, HV22Gs tended to exhibit reduced mRNA levels (Fig. 4g, Supp Fig. 8g-h). 22G-RNAs that are produced downstream of piRNA targeting establish a positive feedback loop by stimulating further production of 22G-RNAs through RNA dependent RNA polymerase (Sapetschnig et al., 2015), but also promote silencing of their targets, leading to a fine balance between silencing and maintenance of 22G-RNA populations. Coupled with low availability of templates for amplification, this might lead to fluctuations in small RNA levels at targets of silencing pathways. Since 22G-RNAs are heritable (Ashe et al., 2012; Buckley et al., 2012; Luteijn et al., 2012; Shirayama et al., 2012), we propose that these fluctuations manifest as epimutations that can exhibit bistability but last only a few generations (Fig. 4h). In contrast, CSR-1 targets have higher mRNA levels (Supp. Fig. 8i) thus they have more stable 22G-RNA levels and are less liable to epimutations.

**Figure 4.**
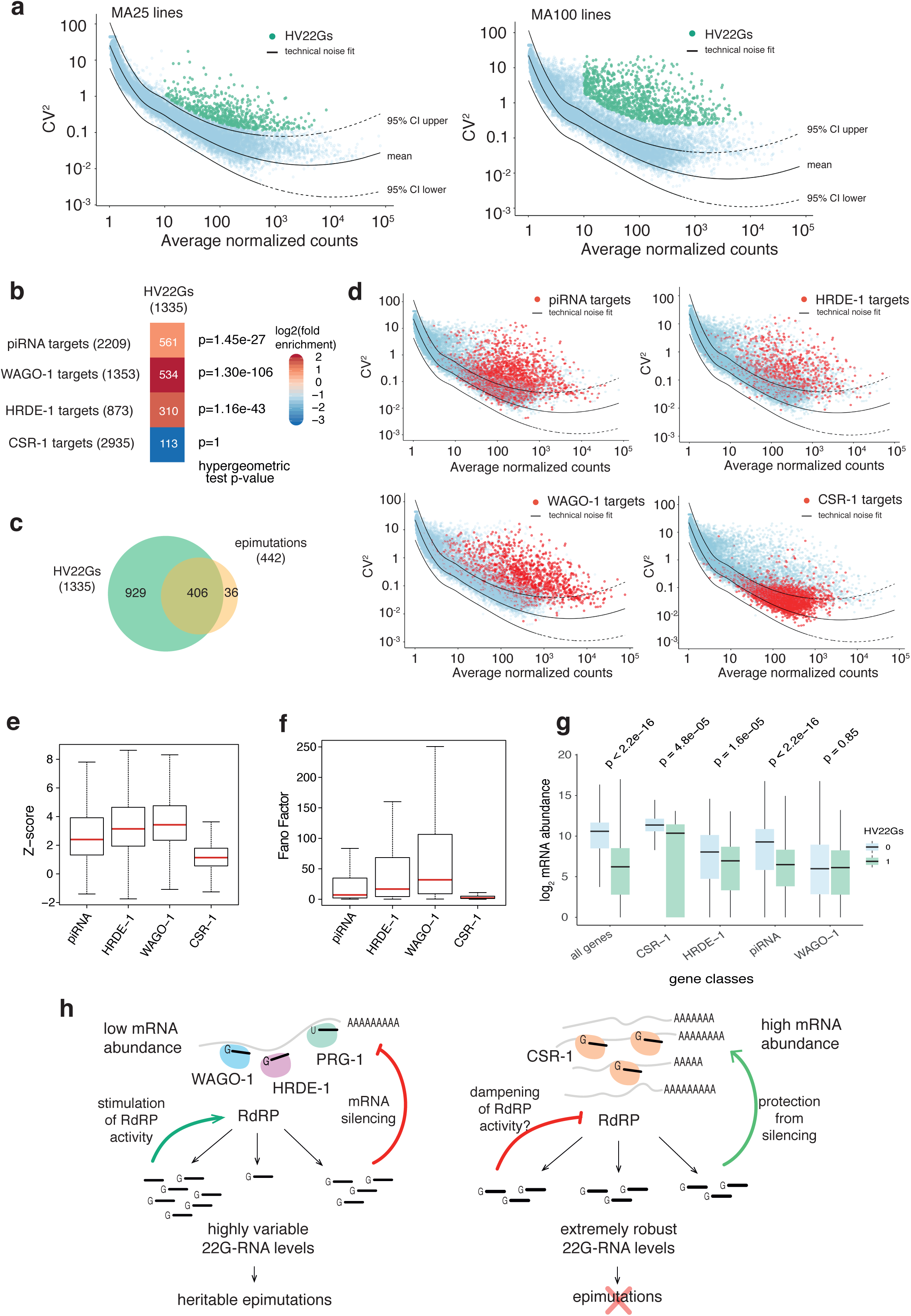
Silencing small RNA pathways show hypervariability in 22G-RNA levels. a. Identification of genes with highly variable 22Gs (HV22Gs). The squared coefficients of variation in 22G-RNA levels are plotted against the mean. The technical noise fit is shown in pink, dashed lines represent 95% confidence intervals for the fit. Genes showing increased variability compared to technical noise are highlighted in red. b. Overlap of HV22Gs with small RNA pathway target genes. The significance of the overlap is represented by the colour of the heatmap. c. Overlap of HV22Gs with epimutable genes. d. Comparison of variability in 22G-RNA levels for different small RNA pathway gene targets. e-f. Quantification of variability in 22G-RNA levels for different small RNA pathway gene targets. E shows Z-score distributions calculated on the basis of the distribution of technical noise for each of the gene classes. F shows the distribution of Fano factors for each of the gene classes. Box plot shows interquartile range with a line at median, and whiskers extend to the greatest point no more than 1.5 times the interquartile range. g. Comparison of mRNA abundance of HV22Gs and non-HV22Gs for all genes, and within small RNA pathway gene targets. Genes with >20 normalised 22G counts were considered for this analysis. Box plot shows interquartile range with a line at median, and whiskers extend to the greatest point no more than 1.5 times the interquartile range h. Model for the emergence of epimutations. CSR-1 target genes do not undergo epimutations due to (1) high mRNA abundance and (2) the protective role of CSR-1 from silencing activities. In contrast, silencing small RNA pathway targets show a high level of variability in 22G-RNAs. This arises due to a combination of (1) low mRNA abundance, and (2) the ability of 22G-RNA populations to self-sustain, establishing a positive feedback for 22G-RNA amplification. Epimutations are an extreme example of this process, leading to heritable epigenetic variation.

Taken together, our results provide the first demonstration that small RNA-mediated epimutations arise during experimental evolution in a metazoan organism. We measure their rate, spectrum, and stability and provide a plausible mechanism for their formation and disappearance. Our results suggest that epigenetic inheritance carried by small RNAs and traditional genetic inheritance carried by DNA sequence alterations exist at opposite ends of a spectrum: epimutations arise rapidly but have limited stability, while mutations arise at a lower rate but are stably inherited. Epigenetic inheritance thus offers the potential to stimulate adaptation over short timescales, but is unlikely to contribute to long-term inheritance without contribution from DNA sequence changes. Interestingly, the short-term nature of epimutations driven by small RNAs in *C. elegans* also characterizes epimutations due to DNA methylation in plants (Becker et al., 2011; van der Graaf et al., 2015; Hagmann et al., 2015; Schmitz et al., 2011), and indeed *C. elegans* possesses mechanisms that limit the duration of RNAi silencing (Lev et al., 2017; Perales et al., 2018).

## Supplementary Figure Legends

**Supplementary Figure 1.**
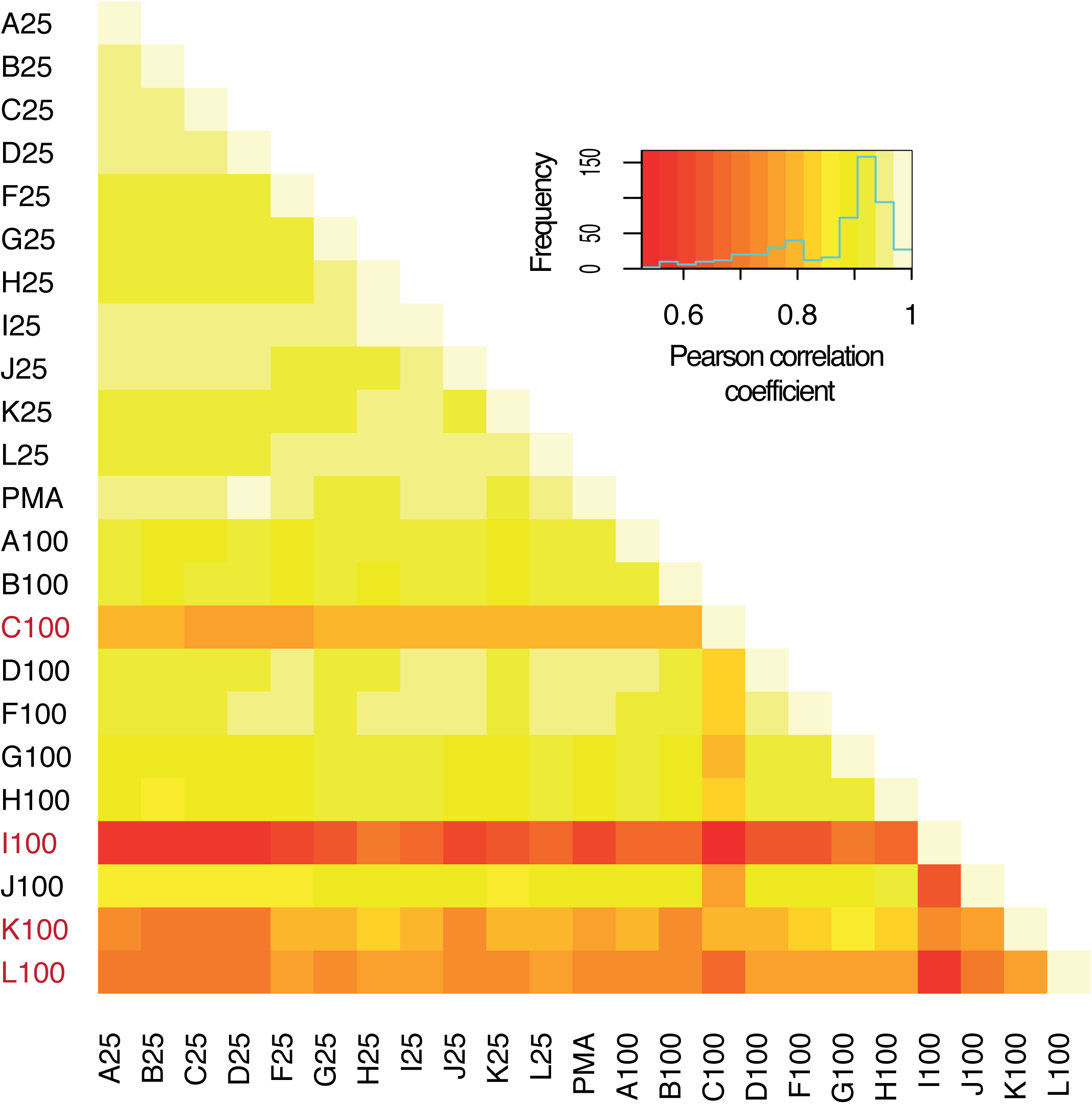
Correlation analysis of the 22G-RNA counts dataset. Heatmap depicting all vs all correlation values calculated for the set of epimutated genes in MA25 and MA100 lines, after log2 transformation of the normalised counts matrix.

**Supplementary Figure 2.**
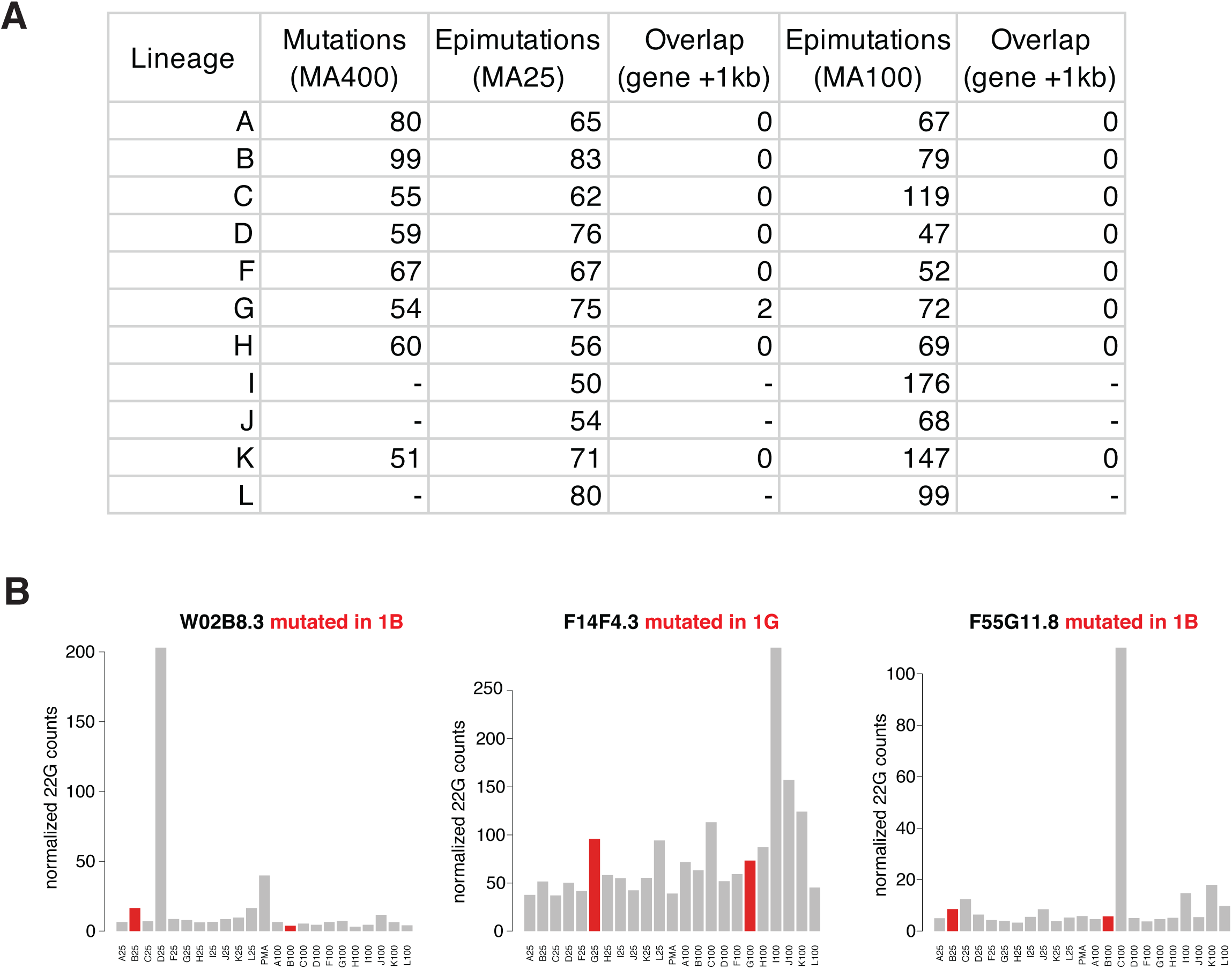
Minimal overlap of epimutations and genetic mutations across lineages. a. Overlap between genetic mutations detected by high throughput genome sequencing after 400 generations of selfing (MA400 lines, Konrad et. al., 2019), and epimutations detected in MA25 and MA100 lines. b. Representative examples of 22G-RNA counts across lines for epimutable genes with an overlapping mutation in any one line, regardless of their epimutation status. A mutation was considered to overlap with a gene if located within the gene or 1kb flanking regions. Lines with an overlapping mutation are shown in red. c. 22G-RNA counts in the two epimutations in lineage G overlapping with genetic mutations as in b.

**Supplementary Figure 3.**
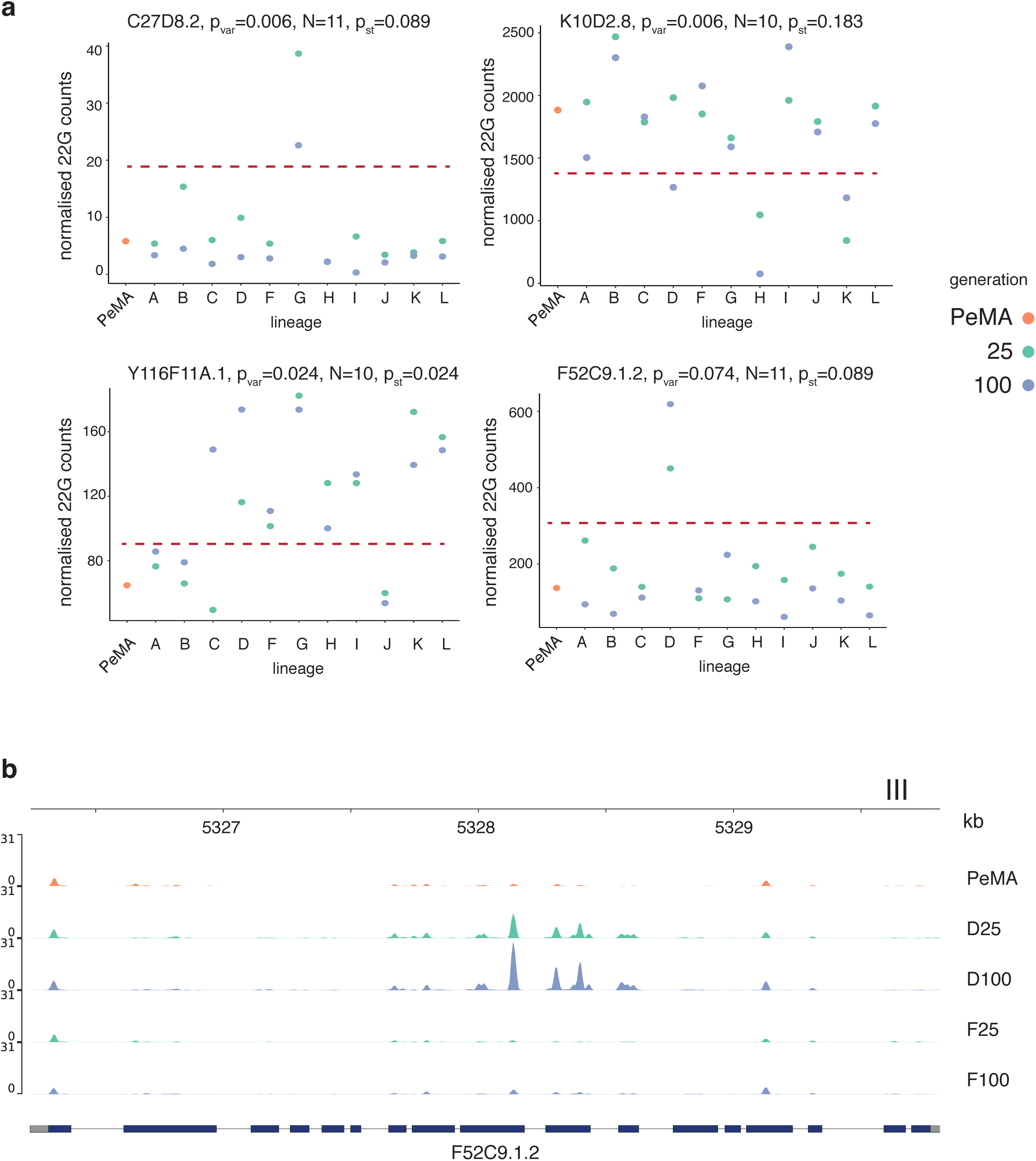
Examples of long-lasting epimutations and genes with reduced within-lineage variation. a. 22G normalised counts across lines for genes showing low p-values in both the variance test (p_var_) and the epistates test (p_st_). N indicates the number of matching states. The red line indicates the threshold separating high and low small RNA states according to k-means clustering. b. 22G signal profile for F52C9.1.2 in the parental line (PeMA), in lineage D (epimutated) and lineage F (no change in 22G-RNAs).

**Supplementary Figure 4:**
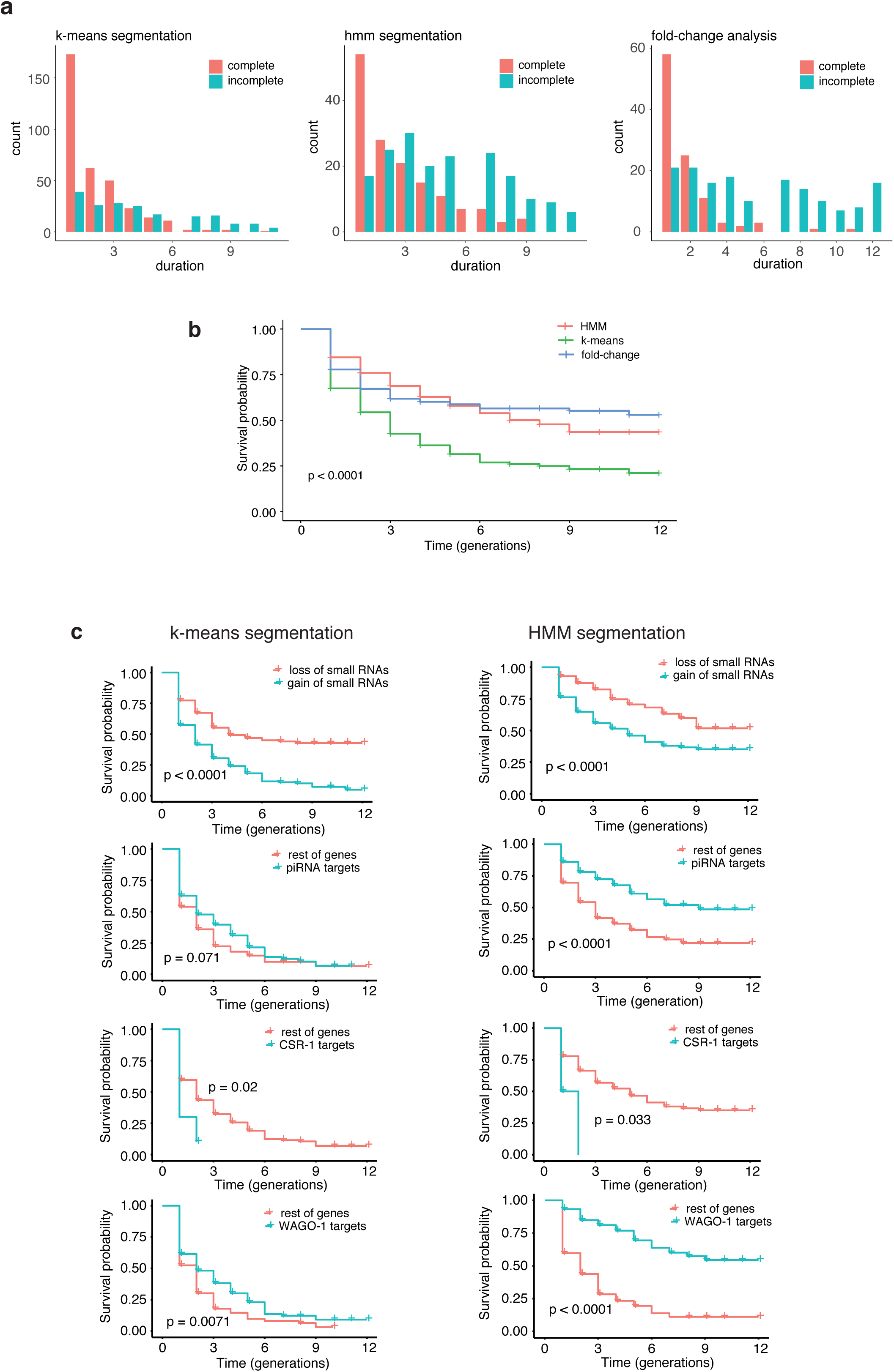
Comparison of methods for survival analysis of epimutations. a-b. Comparison of methods to estimate the duration of epimutations. Duration distributions are shown in A, and survival curves in B. c. Genomic features influencing the duration of epimutations. Survival curves estimated from either the k-means or the HMM datasets are shown, showing qualitatively similar trends.

**Supplementary Figure 5:**
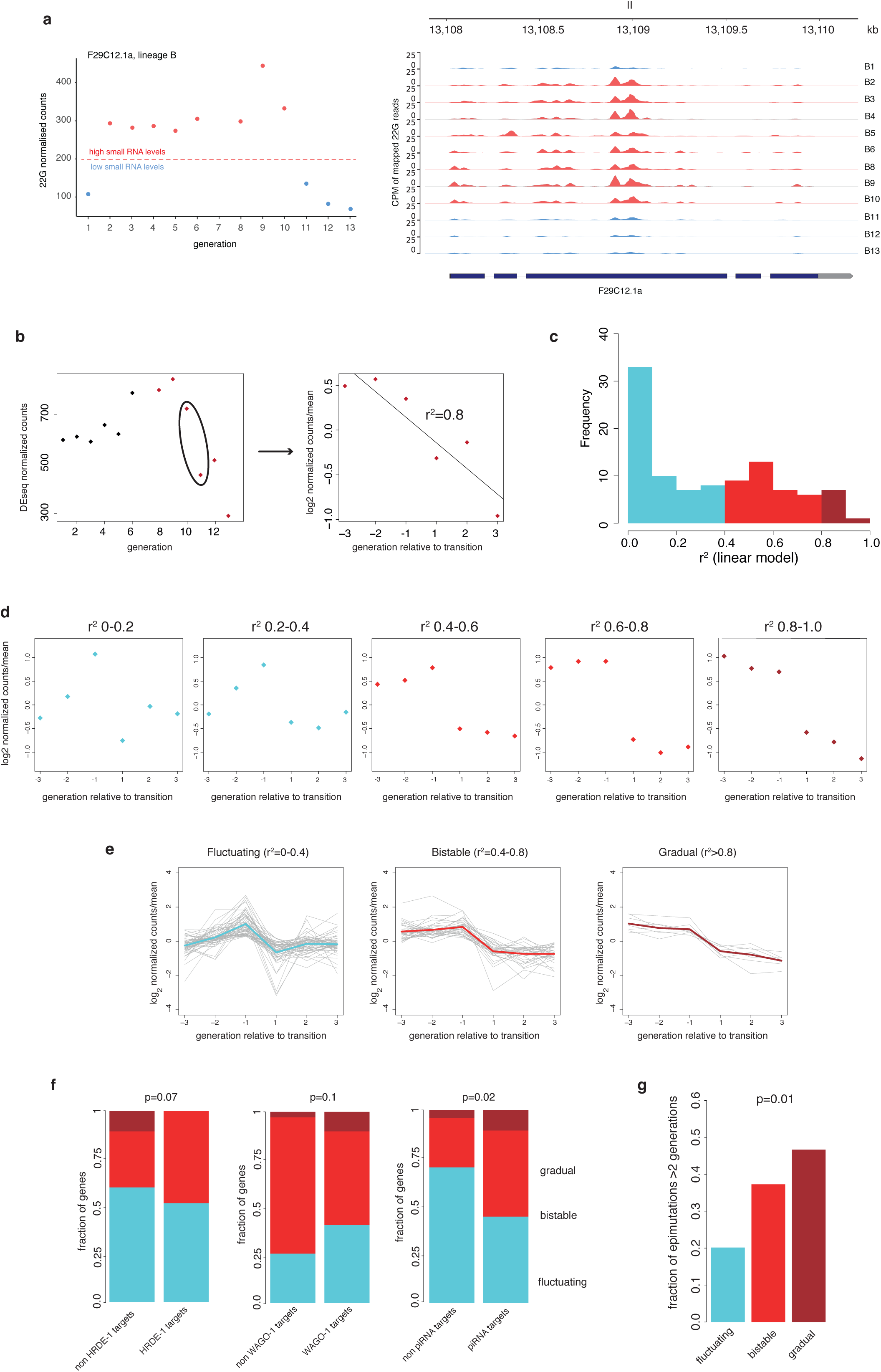
Dynamics of transgenerational 22G-RNA changes. a. Example of a gene displaying bistable 22G-RNA levels (compare with Fig. 3d). b. Method to examine the transition in 22G-RNA levels around the epimutation. Three generations before and after the transition were selected, and a linear model was fit to the data, recovering an r^2^ that represents the linearity of the change. c. Histogram illustrating the distribution of r^2^ values obtained from the analysis in b applied to all epimutations. d. Mean normalized small RNA levels within groups spanning different linearity in 22G-RNA alterations. e. Classification into fluctuating, bistable and gradual changes in small RNAs. Grey lines show each individual epimutation and the mean across each group is shown as a thick coloured line. f. Association of different 22G-RNA dynamics with HRDE-1, piRNA and WAGO-1 target genes. The p-value for a difference in proportion between targets and non-targets according to a Fisher’s exact test is shown above each plot. g. The duration of bistable and gradual changes is longer than fluctuating epimutable genes. The p-value for a difference in proportion between <=2 or >2 generations of epimutations is shown above the plot.

**Supplementary Figure 6:**
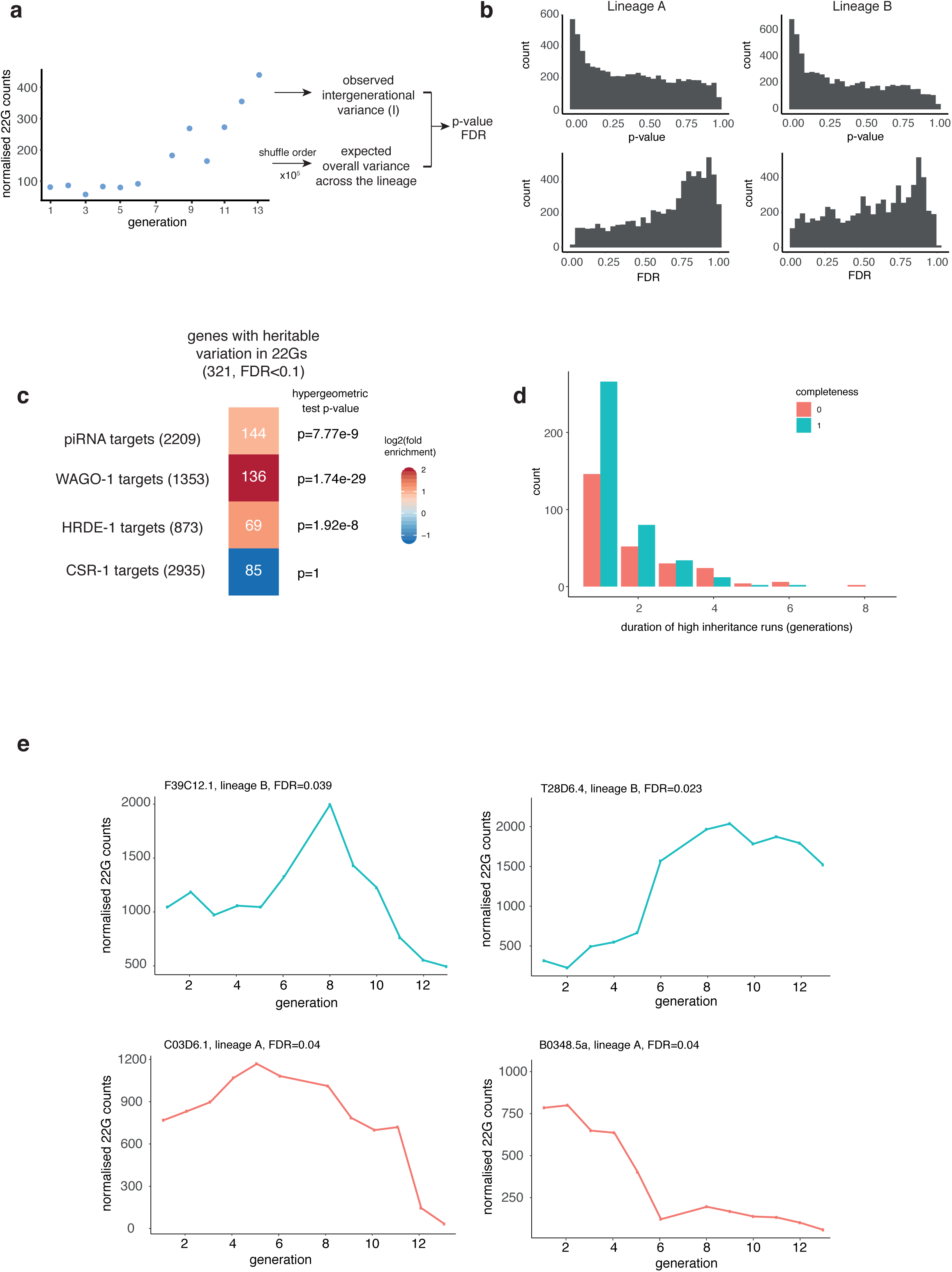
Direct identification of genes with heritable variation in 22G-RNA levels. a. Test for short-term inheritance. By comparing the intergenerational variance with the overall variance across the lineage, genes with heritable variation in 22G-RNA levels are identified. b. p-value and FDR histograms from the test for short-term inheritance, in lineages A and B. c. Overlap of genes with heritable 22G-RNA levels with small RNA pathway gene targets. d. Distribution duration of runs of consecutive generations with low intergenerational variance. These were defined as consecutive generations with a difference lower than 30% of the standard deviation in 22G-RNA levels across the lineage. e. Examples of 22G-RNA dynamics at genes with heritable variation in 22G-RNA levels in lineages A and B.

**Supplementary Figure 7.**
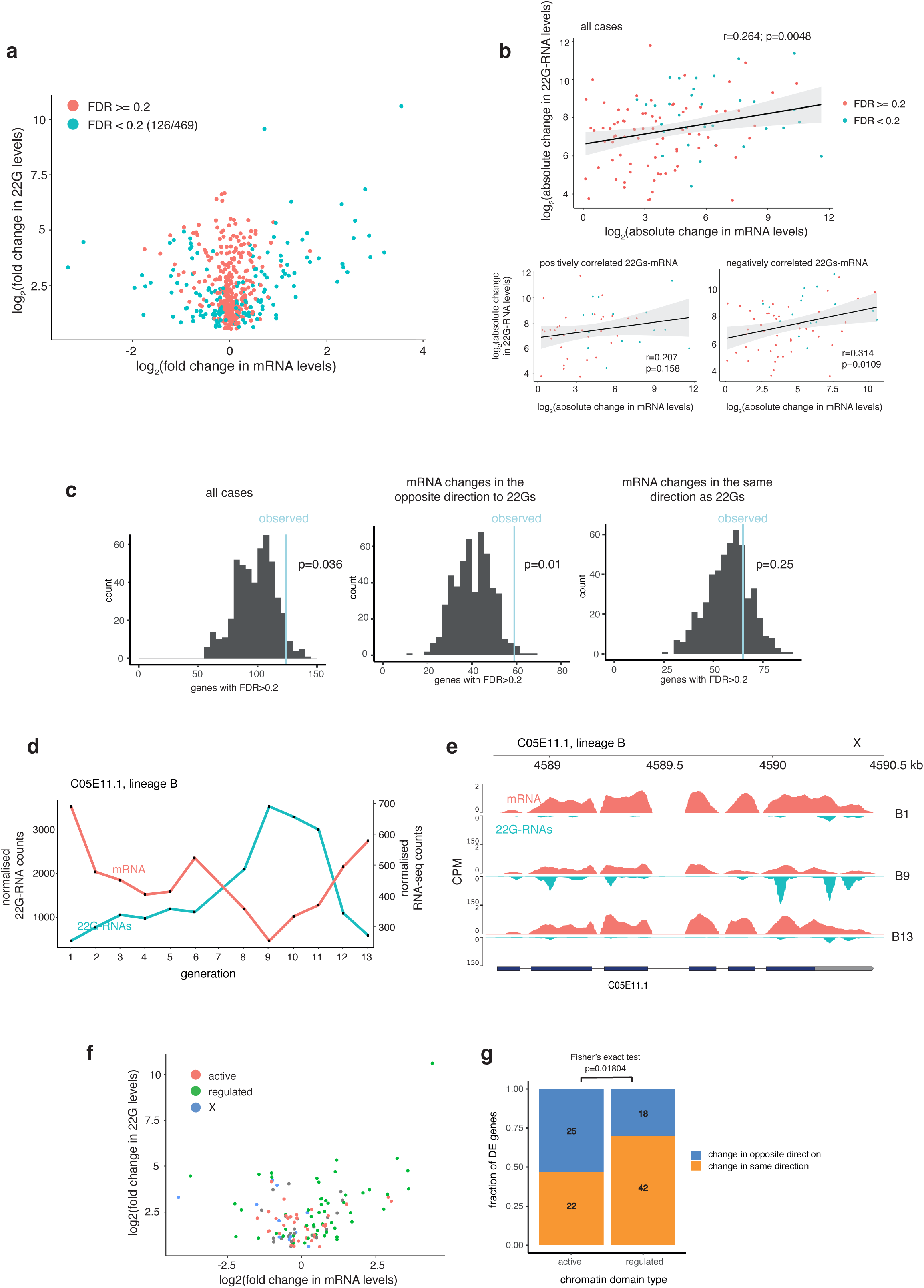
Integration of 22G-RNA and mRNA count data. a. Visualisation of changes in 22G-RNA levels against changes at the mRNA level. Genes with significantly different mRNA levels between the high and low small RNA states are shown in green. b. Correlation between the absolute change in 22G-RNA and mRNA levels, for all epimutations (top panel), and for genes with positively correlated (bottom left) and negatively correlated (bottom right) mRNA-22G changes. c. Genes with correlated changes in 22G-RNAs and mRNAs are marginally enriched in the epimutations set (blue line) compared to random subsets of genes with >10 22G-RNA normalised counts. This enrichment is highest for cases where there is a negative correlation between 22G-RNA and mRNA levels. d-e. Example of gene with negatively correlated 22G-RNA and mRNA levels. f. Visualisation of changes in 22G-RNA levels against changes at the mRNA level for genes with significantly different mRNA levels between the high and low small RNA states, coloured by chromatin domain location. g. Comparison of the proportions of genes with positively and negatively correlated changes in 22G-RNAs and mRNAs in active and regulated chromatin domains.

**Supplementary Figure 8.**
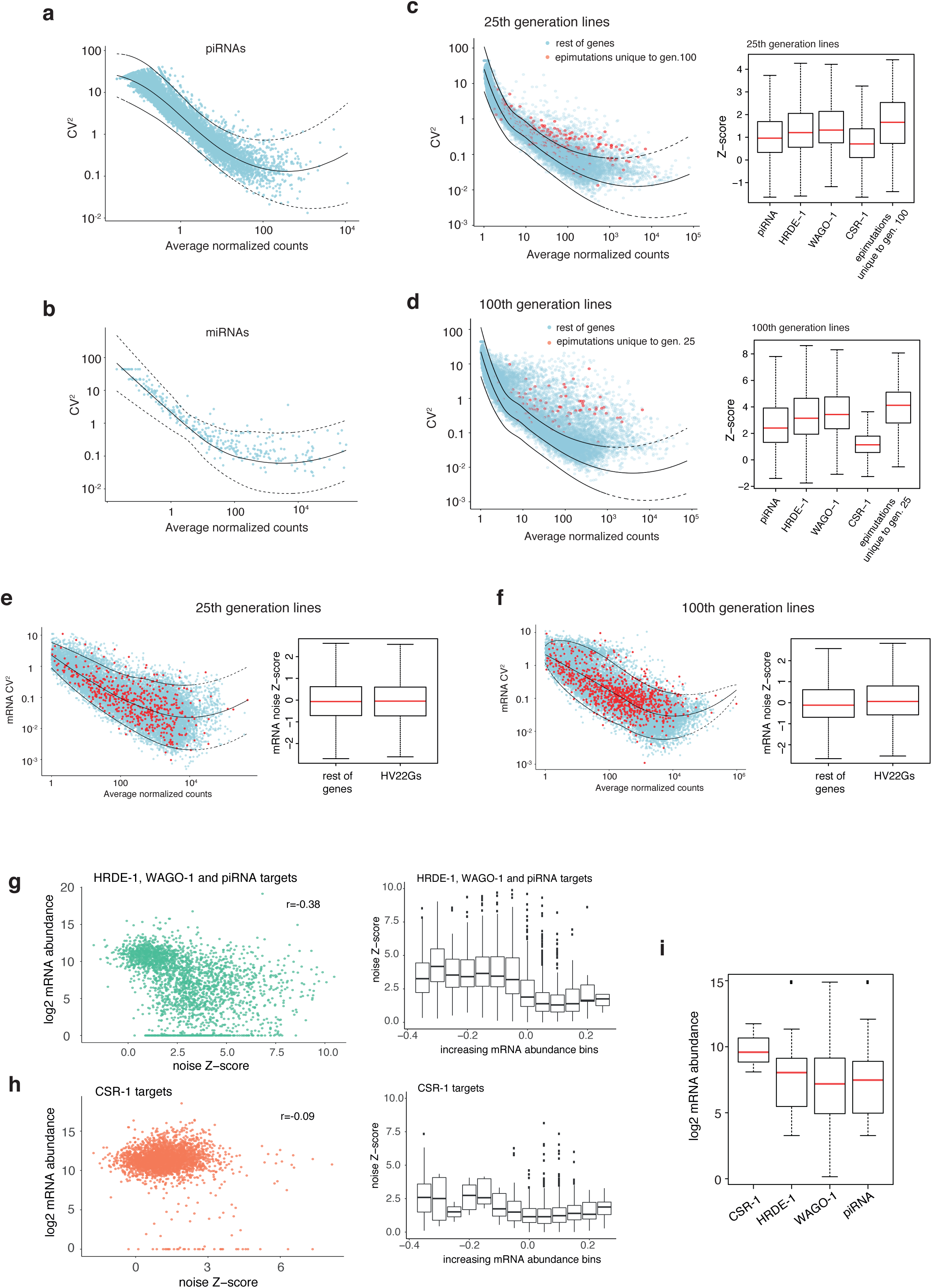
Further analysis of variability in small RNA pathways and their target genes. a-b. Variability analysis of piRNAs (A) and miRNAs (B). c. Epimutations unique to MA100 lines are hypervariable in the MA25 dataset. d. Epimutations unique to MA25 lines are hypervariable in the MA100 dataset. e. Epimutations from MA25 lines do not show increased variation at the mRNA level. f. Epimutations from MA100 lines do not show increased variation at the mRNA level. g-h. Correlation analysis between mRNA abundance and variability in 22G-RNAs, for genes with >20 22G normalised counts, separating piRNA, HRDE-1 and WAGO-1 targets (F) and CSR-1 targets (G). i. Epimutated CSR-1 targets show higher expression than the rest of epimutated genes, consistent with CSR-1 target status.

**Supplementary Figure 9.**
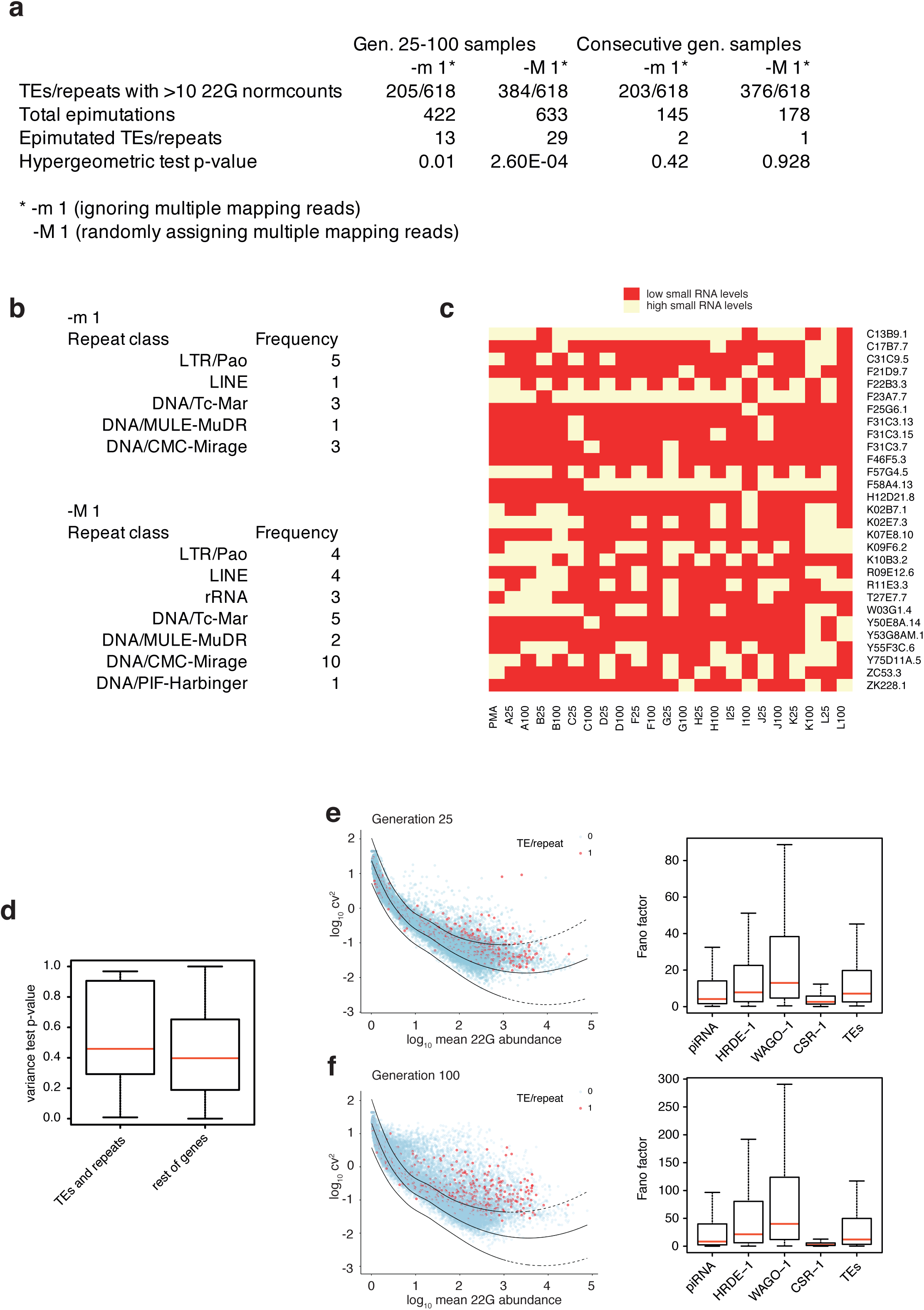
Epimutations at transposable elements and other repeats. a. Summary of epimutations at transposable elements (TEs) and other repeats. TEs are only moderately enriched in the epimutation sets from the MA25 and MA100 lines, and are not enriched in the set of epimutations observed in the consecutive generation experiment. b. Repeat class annotations of epimutated TEs and other repeats. c. Small RNA level states for epimutated TEs and other repeats in MA25 and MA100 lines, showing little correspondence between states. d. Distribution of variance test p-values for TEs and other repeats compared to the rest of genes. No TEs or other repeats had significantly reduced variance within lineages after multiple testing correction. e-f. Variability analysis of 22Gs mapping to TEs and other repeats in MA25 (e) and MA100 (f) lines, showing increased variability in comparison to CSR-1 targets.

## Methods

### Nematode culture

*Caenorhabditis elegans* nematodes were grown in NGM plates seeded with OP50 *E. coli* at 20°C. 25^th^ generation and 100^th^ generation lines were grown as described in ^1^. For the consecutive generation experiment, each lineage was founded by picking a single N2 L4 hermaphrodite worm (Day 1). On Day 4, a single L4 was bottlenecked into a new plate as a founder for the next generation. On Day 5, the rest of the worms were synchronised by hypochlorite treatment, and grown to the young adult stage in order to obtain RNA samples. This procedure was carried out for 13 generations, with exception of generation 7 where the RNA sample collection was not carried out.

### Preparation of RNA samples and small RNA libraries

Synchronised young adult hermaphrodites were washed off plates using M9, and collected into 15mL tubes. Worms were washed 3 additional times with M9, and 1mL Trizol was added for each 100*µ*l of worms to proceed with Trizol-chloroform RNA extraction. RNA was precipitated overnight by adding 1*µ*l glycogen and an equal volume of isopropanol.

For small RNA library preparation, 1*µ*g (lineage A and B samples) or 2*µ*g (25^th^ and 100^th^ generation samples) of RNA were incubated for 1h at 37°C with 5’ RNA polyphosphatase (Epicentre) at a final concentration of 1U/*µ*l, in a total volume of 20ul, in order to convert 5’PPP 22G-RNAs into clonable 5’P 22G-RNAs. RNA was phenol-chloroform extracted and precipitated overnight with 1*µ*l glycogen, 1/10 volume 3M AcNA and 3 volumes 100% ethanol. Small RNA libraries were generated using the Illumina TruSeq Small RNA library preparation kit as per manufacturer’s instructions, and gel-purified to size select 21-23nt inserts. For mRNA-seq, 1*µ*g of total RNA was spiked-in with 100ng of *Schizosaccharomyces pombe* total RNA, and the mix was subjected to poly-A selection using the NEBNext polyA Magnetic Isolation Module. RNA-seq libraries were prepared using the NEBNext Ultra II Directional RNA Library Prep Kit for Illumina. Both types of libraries were sequenced on an Illumina HiSeq2500 instrument.

### Processing of small RNA sequencing and RNA-seq data

Small RNA libraries were trimmed using cutadapt v1.10^2^, and reads >18nt and <35nt were mapped using Bowtie v0.1.2^3^ with parameters -v -m1 to the WS252 *C. elegans* genome. 22G reads were mapped with the same parameters to a *C. elegans* transcriptome file including WS252 annotated mRNA transcripts, ncRNA transcripts, pseudogenic transcripts, and transposon transcripts. For genes with multiple RNA isoforms, the longest isoform was selected and the rest of isoforms were filtered out. Antisense 22G read counts were computed for each RNA in the reference transcriptome. To visualise the signal of 22G-RNAs across the genome, bam files were converted to a bigwig format using deeptools v3.1.2^4^ bamCoverage with parameters -bs 10 --smoothLength 30 --normalizeUsing CPM.

Additionally, we mapped the small RNA reads with parameters -v 0 -M 1 to randomly assign multiple mapping reads in order to capture changes in 22G-RNAs derived from repetitive regions of the transcriptome, and we analyzed the two datasets (-m 1 and -M 1) in parallel.

RNA-seq reads were mapped using Tophat v2.0.11^5^ with parameters -i 30 -I 20000, to a combined reference index containing the WS252 *C. elegans* genome and the *Schizosaccharomyces pombe* genome (2018-06-01 PomBase release). *C. elegans* counts were recovered using htseq-count 0.9.1^6^, and a gtf file including longest isoform RNA annotations as described above. *S. pombe* counts were recovered using htseq-count, and the corresponding gtf file from the 2018-06-01 PomBase release.

### Identification of epimutations

RNA-seq and 22G-RNA counts were normalised using DEseq2^7^. Epimutations between any two samples were identified using the 22G-RNA data. log2(fold changes) were plotted against the log2(mean counts) for each gene (MA-plot), and a loess smoothing curve was fitted to the scatterplot. For each data point, a Z-score was derived by calculating the difference of the observed fold change minus the average of the loess fit, divided by the standard deviation of the loess fit. P-values were derived using a normal distribution, and corrected for multiple testing using the Benjamini-Hochberg False Discovery Rate (FDR) method^8^.

We defined epimutable genes in the 25^th^ and 100^th^ generation samples as those below an FDR threshold of 1e-4 in at least one pairwise comparison. Correlation analysis was done on this subset of genes, after log2 transforming the count matrix+1 pseudocount. This analysis revealed that C100, I100, K100 and L100 have accumulated large differences in 22G levels in a large number of genes and are outliers compared to all the rest of samples. On the basis of this observation, we performed the subsequent analyses both in the absence and presence of these samples separately.

We defined epimutable genes in the consecutive generation samples as those with an FDR threshold of 1e-4 in at least one pairwise comparison. Correlation analysis was done on this subset of genes, after log2 transforming the count matrix+1 pseudocount. This analysis revealed two groups of samples corresponding to early (1-8) and late (9-13) in the lineage. Since this effect likely corresponds to experimental variation, genes that showing systematically different levels of 22Gs between the two groups of samples as detected by DEseq2 (FDR<0.1) were excluded from the downstream analysis. Samples B1 and B4 behaved as outliers compared to the rest of samples, and, similarly, genes with different levels of 22Gs in these two samples were excluded from further analysis.

For each gene, we used unsupervised k-means clustering on the normalised counts for all samples, including those from the consecutive generations experiment. This allowed us to classify the samples in high and low small RNA level groups. The total number of epimutations between any two samples was calculated on the basis of the group assignations.

### Test for long-term epigenetic inheritance

For each gene, we calculated the number of matches (N) in the small RNA level states (high or low) between generation 25 and their corresponding generation 100 samples. We then randomised those states in both lineages 10^5^ times in order to obtain a null distribution of N. A p-value was calculated as the fraction of cases in the null distribution where the simulated N equals or exceeds the observed N. In addition, in order to incorporate the quantitative range of the data into the test, we calculated for each gene the observed within-lineage variance (W) using the normalised count data, as the sum of the squared difference in 22G counts between samples of matching lineages, divided by the total number of lineages (11):

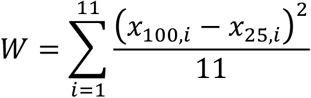

We then randomised the count data in both lineages 10^5^ times in order to obtain a null distribution of W. A p-value was calculated as the fraction of cases in the null distribution where the simulated W is equal or less than the observed N. In both tests, p-values were corrected for multiple testing using the FDR method.

### Test for short-term epigenetic inheritance

For each gene, we calculated an intergenerational variance (I) considering only pairs of consecutive generations:

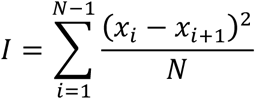

We then randomised the order of the data points 10^5^ times in order to obtain a null distribution of I, that reflects the overall variance across the lineage. A p-value was calculated as the fraction of cases in the null distribution where the simulated I is equal or less than the observed I. p-values were corrected for multiple testing using the FDR method.

Using the set of genes with significantly low intergenerational variance, we defined pairs of consecutive generations as showing high levels of inheritance if the difference in small RNA levels was less than 30% of the standard deviation of the set of data points across the lineage. We then quantified the distribution of durations of runs of high inheritance across all genes. The median duration of high inheritance runs was derived using survival analysis, incorporating censoring information to reflect that fact that some runs reach the end of the lineage, such that their exact duration is unknown.

### Quantification of epimutation duration

We quantified epimutation duration using three different methods. Firstly, we used the high and low small RNA state assignations across the lineage for the set of epimutable genes. Secondly, we used depmixS4^9^ to fit a 2-state Hidden Markov model to the count data across the lineage, in order to recover the trajectory of hidden states corresponding to high and low small RNA levels. The HMM fit was initialised by providing the mean of the high and low small RNA level states as initial guesses. Finally, in order to incorporate the continuous nature of the data more directly, we identified pairs of consecutive generations with a fold-change greater than 2 after adding 10 pseudocounts to the normalised 22G counts. For each of these, we calculated the number of generations for which the difference in small RNA levels remains equal or greater than 2-fold. We derived from these a distribution of epimutation duration, considering complete (finished within the lineage) and incomplete (reaching the end of the lineage) epimutations separately.

We used survival analysis to quantify the overall stability of epimutations, incorporating the duration of complete and incomplete epimutations, considering incomplete epimutations as censored observations. We tested the effect of 22G fold changes and mRNA fold changes on epimutation duration using a Cox proportional hazards model. We tested for differences in epimutation duration depending on small RNA pathway regulation using Kaplan-Meier estimators and a log-rank test.

To investigate the dynamics of the transitions between different small RNA states at epimutated genes we extracted 3 generations either side of the largest difference between successive generations to give a time period of 6 generations in total. Only genes where 3 generations either side of the transition could be extracted were considered. We normalized to the mean across the 6 generations and used a linear model, log(x)=a.(t)+b where x is normalized 22G-RNAs, t is generations relative to the transition, i.e. -2<=t<= 3, and a and b are constants. The r^2^ was extracted from the fit. Qualitative examination of small RNA dynamics with different r^2^ values suggested three categories, which we labelled as fluctuating (r^2^ <0.4), bistable (0.4<= r^2^ <0.8) and gradual (r^2^ >=0.8), and these were used for further intersection with different categories of genes (see below for definition of gene classes).

### Integration of RNA-seq and small RNA sequencing data

To analyze changes in mRNA expression associated with changes in 22G-RNA levels, we used a Wilcoxon rank sum test to compare the RNA-seq normalised counts between groups of samples with high and low small RNA levels, defined by k-means clustering as described above. We applied this analysis to all epimutable genes, using the 25^th^ and 100^th^ generation samples, and the consecutive generation samples. Multiple testing correction was applied using the Benjamini-Hochberg FDR method. We examined the relationship between the absolute change in 22G-RNA levels and mRNA levels, showing significant positive correlations.

To test whether genes with significant changes in mRNA levels between high and low small RNA level samples are enriched in the set of epimutable genes, we sampled 422 genes from the entire set of genes with >10 normalised counts times, calculated high and low 22G-RNA level groups, and determined the number of genes from this set also showing significant changes in mRNA levels as described above. We repeated this analysis 10,000 times to generate a null distribution of the expected number of genes showing significant changes in mRNA levels, and compared it to the observed value in the set of epimutable genes. We examined all cases, and cases with positively and negative correlated 22G-RNA and mRNA levels separately.

### Variability analysis and identification of genes with hypervariable 22Gs (HV22Gs)

We used the 25^th^ and 100^th^ generation samples to quantify the variation in 22G-RNA levels across lineages transcriptome-wide, and identify HV22Gs. To do this, we estimated the technical variance as the sum of squared differences between pairs (*x*_*i*_, *y*_*i*_) of technical duplicates of libraries, divided by 11 (the total number of pairs).

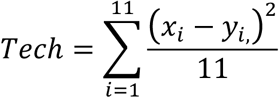

We plotted the technical coefficients of variation against the mean 22G levels in a logarithmic scale, showing a decreasing relationship as typically seen in mRNA sequencing data. We then fit a loess smoothing curve, and used this fit as a baseline to identify HV22Gs.

We estimated the total observed variance as the sum of squared differences between all pairs of libraries, divided by the total number of comparisons:

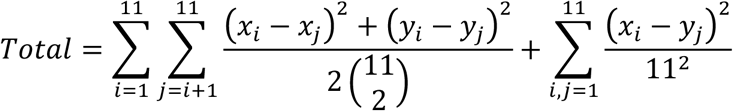

We plotted the total coefficient of variation against the mean 22G levels in a logarithmic scale, together with the technical variation fit (mean and 95% confidence intervals). In order to identify HV22Gs, we calculated the residuals of the total variance data to the technical fit, and divided by the standard deviation of the technical fit to obtain a Z-score and a p-value based on a normal distribution. We corrected for multiple testing using the FDR method.

To estimate the technical variation in mRNA-seq data, we used the counts from the *S. pombe* spiked-in total RNA. To remove variation due to spike-in ratios and sequencing depth, we downsampled each library to the minimum observed number of *S. pombe* and *C. elegans* total reads within (for the 25^th^ and 100^th^ generation data separately). We used two baselines to quantify mRNA variability, (1) the technical variation estimated from *S. pombe* counts, and (2) the overall variation observed from *C. elegans* counts.

### Definition of gene classes

We sourced publicly available small RNA sequencing data from IP experiments for HRDE-1 (GSM948684 and GSM984685), WAGO-1 (SRR030711 and SRR030712) and CSR-1 (SRR030720 and SRR030721), processed it as described above, and defined their target genes as genes with at least 10rpm in the IP sample, and a 1.5-fold enrichment in the IP sample. piRNA targets were defined as genes exhibiting at least a two-fold decrease in 22Gs in a prg-1 background (SRR2140760 and SRR2140763), and 20rpm of 22Gs in the WT. Chromatin domain assignations were taken from Evans et. al., 2016 and lifted over to WS252. To annotate repetitive transcripts, we ran RepeatMasker v4.0.5^10^ with parameters -nolow on the transcriptome file. Repetitive transcripts were defined as those with at least 80% of their length covered by RepeatMasker hits.

## Data accessibility

Small RNA sequencing and RNA-seq datasets have been deposited in the Sequence Read Archive (PRJNA553063).

